# Capillary Morphogenesis gene 2 mediates multiple pathways of growth factor-induced angiogenesis by regulating endothelial cell chemotaxis

**DOI:** 10.1101/705442

**Authors:** Lorna Cryan, Tsz-Ming Tsang, Jessica Stiles, Lauren Bazinet, Sai Lun Lee, Samuel Garrard, Cody Roberts, Jessie Payne, P. Christine Ackroyd, Kenneth A. Christensen, Michael S. Rogers

## Abstract

Pathological angiogenesis contributes to diseases as varied as cancer and corneal neovascularization. The vascular endothelial growth factor (VEGF) - VEGF receptor 2 (KDR/VEGFR2) axis has been the major target for treating pathological angiogenesis. However, VEGF-targeted therapies exhibit reduced efficacy over time, indicating that new therapeutic strategies are needed. Therefore, identifying new targets that mediate angiogenesis is of great importance. Here, we report that one of the anthrax toxin receptors, capillary morphogenesis gene 2 (ANTXR2/CMG2), plays an important role in mediating angiogenesis induced by both bFGF and VEGF. Inhibiting physiological ligand binding to CMG2 results in significant reduction of corneal neovascularization, endothelial tube formation and cell migration. We also report the novel finding that CMG2 mediates angiogenesis by regulating the direction of endothelial chemotactic migration without affecting overall cell motility.

## Introduction

Pathological angiogenesis plays a role in many diseases, including cancers and various ocular diseases that lead to blindness. Most of the current treatment strategies are targeting the VEGF - VEGFR2 axis. However, such therapies only provide modest efficacy overtime and are accompanied unpleasant side effects. Thus, identification of new anti-angiogenic targets are needed and critical for development of alternative therapeutic strategies. It was previously demonstrated that one of the anthrax toxin receptors, CMG2, has a role in mediating angiogenesis^1, 2^. Our previous work demonstrated that inhibition of CMG2 potently reduces angiogenesis in the cornea^1^ and inhibit endothelial cell migration^3, 4^. Thus, we are interested to investigate the role(s) of CMG2 in mediating angiogenesis.

While knowledge of CMG2’s exact role in angiogenesis remains limited, CMG2’s function as one of anthrax toxin receptors is very well established. CMG2 and its homolog, tumor endothelial marker 8 (ANTXR1 / TEM8), are the primary sites of anthrax toxin entry into the cells. CMG2 and TEM8 share 40% amino acid homology, including homology in an intracellular domain of unknown function that is shared with no other protein in the mammalian genome^5^. Within their cell-surface von Willebrand Factor A (VWA) ligand-binding domains, homology between CMG2 and TEM8 rises to 60%^6^. During anthrax intoxication, the 83kDa protective antigen (PA), one of three protein subunits from *Bacillus anthracis*, binds to the VWA ligand binding domain of the receptor. PA is then cleaved by a furin-like protease that cuts and releases a 20kDa fragment, leaving a 63kDa PA at the receptor surface. The cleaved PA oligomerizes to form a PA-receptor heptamer, which acts as a binding platform for the other two toxin subunits, lethal factor (LF) and edema factor (EF), Together, these toxin subunits form the complete anthrax toxin, which is then trafficked into the cell with via clathrin-mediated endocytosis^7^.

The endogenous functions of the anthrax toxin receptors in the absence of anthrax toxin are still poorly understood, although, it has been suggested that these receptors interact with extra-cellular matrix (ECM) proteins^2, 8^. In particular, a series of studies have observed observed that mutations on CMG2 or TEM8 ead to build-up of hyaline materials that result in alteration of of skeletal growth and/or alternation of vascular patterns. For example, Loss-of-function mutations in TEM8 cause GAPO syndrome, a disease characterized by vascular anomalies, skeletal defects, growth retardation and progressive fibrosis of various organs^9, 10^. Mutations in CMG2 in humans cause Hyaline Fibromatosis Syndrome (HFS), a rare but serious autosomal recessive disorder that is characterized by accumulation of hyaline material in connective tissue in the skin and other organs and by the presence of non-cancerous nodules^11–,13^ containing excess collagen I, collagen VI, and glycosaminoglycans. To explain the symptoms of HFS, it has been hypothesized that patients with a CMG2 mutation have a defect in either the synthesis or degradation of ECM^12, 14^, presumably related to CMG2 dysfunction.

In addition to ECM binding and trafficking anthrax toxins, it has been suggested that ANTXRs have angiogenic related functions^1, 15^. Our previous work demonstrated that a furin-cleavage-site mutant of protective antigen (PA^SSSR^) inhibits basic fibroblast growth factor (bFGF)-induced corneal neovascularization, VEGF-induced corneal neovascularization, and tumor growth^1, 16^. However, because PA^SSSR^ is known to bind both anthrax toxin receptors (CMG2 and TEM8) as well as integrin β1^17, 18^, which of these receptors was responsible for mediating the observed anti-angiogenic effect of PA^SSSR^ *in vivo* was not firmly established. CMG2 has a much higher affinity for PA than the other two receptors^18–20^ and has been considered the major receptor for PA^21^. Thus, we hypothesized that CMG2 was the likely receptor responsible for the anti-angiogenic effects of PA^SSSR^. Work described here outlines experiments that firmly establish CMG2 as the receptor mediating the observed anti-angionic impact of PA^SSSR^, and provide insight into the role of CMG2 in mediating angiogenesis in endothelial cells.

## Result and Discussion

### Comparison of roles for CMG2 and TEM8 in corneal angiogenesis

As mentioned previously, PA^SSSR^ administration inhibits angiogenesis as measured by endothelial cell migration, corneal neovascularization, and tumor growth^1, 16^. PA^SSSR^ is known to interact with both anthrax toxin receptors and integrin β1^17, 18^. Hence, each of these interactions could, in theory, be responsible for the observed anti-angiogenic effects *in vivo*. CMG2 and TEM8 bind PA^SSSR^ much more tightly than does integrin β1 (CMG2 *K*_d_ = 200pM; TEM8 *K*_d_ = 100nM^18, 22^; integrin β1 *K*_d_ = 1μM), and are therefore more likely to mediate the antiangiogenic effects of PA^SSSR^ than is integrin β1. We performed a series of experiments to compare the relative contributions of CMG2 and TEM8 to corneal neovascularization in mice, and to determine whether blockade of either of these receptors was sufficient to explain the observed anti-angiogenic effects of PA^SSSR^. First, we carried out experiments that used either a CMG2 or TEM8 extracellular domain fused to an antibody Fc-domain to disrupt the ligand-receptor interaction by competing for binding of endogenous ligand. In the corneal micropocket assay, administration of CMG2-Fc significantly inhibited bFGF-induced vessel growth when compared to the untreated control, but the TEM8-Fc fusion did not (Fig. 1A). Hence, targeting CMG2 impacted corneal angiogenesis, while targeting TEM8 did not. We confirmed this observation by administering antibodies specific to either the CMG2 or TEM8 extracellular domains. The presence of the anti-CMG2 antibody significantly reduced bFGF-induced corneal neovascularization in a concentration-dependent manner (Fig. 1B). In contrast, treatment with the anti-TEM8 antibody L2 at a dose and schedule previously shown to inhibit tumor growth in mice (20 mg/kg/week)^23^ resulted in no significant decrease in corneal neovascularization compared to the vehicle control (Fig. 1C). Together, these results show that inhibiting the interaction of CMG2 with its physiological ligand significantly inhibits corneal angiogenesis, while inhibition of the TEM8-ligand interaction does not.

**Figure 1.**
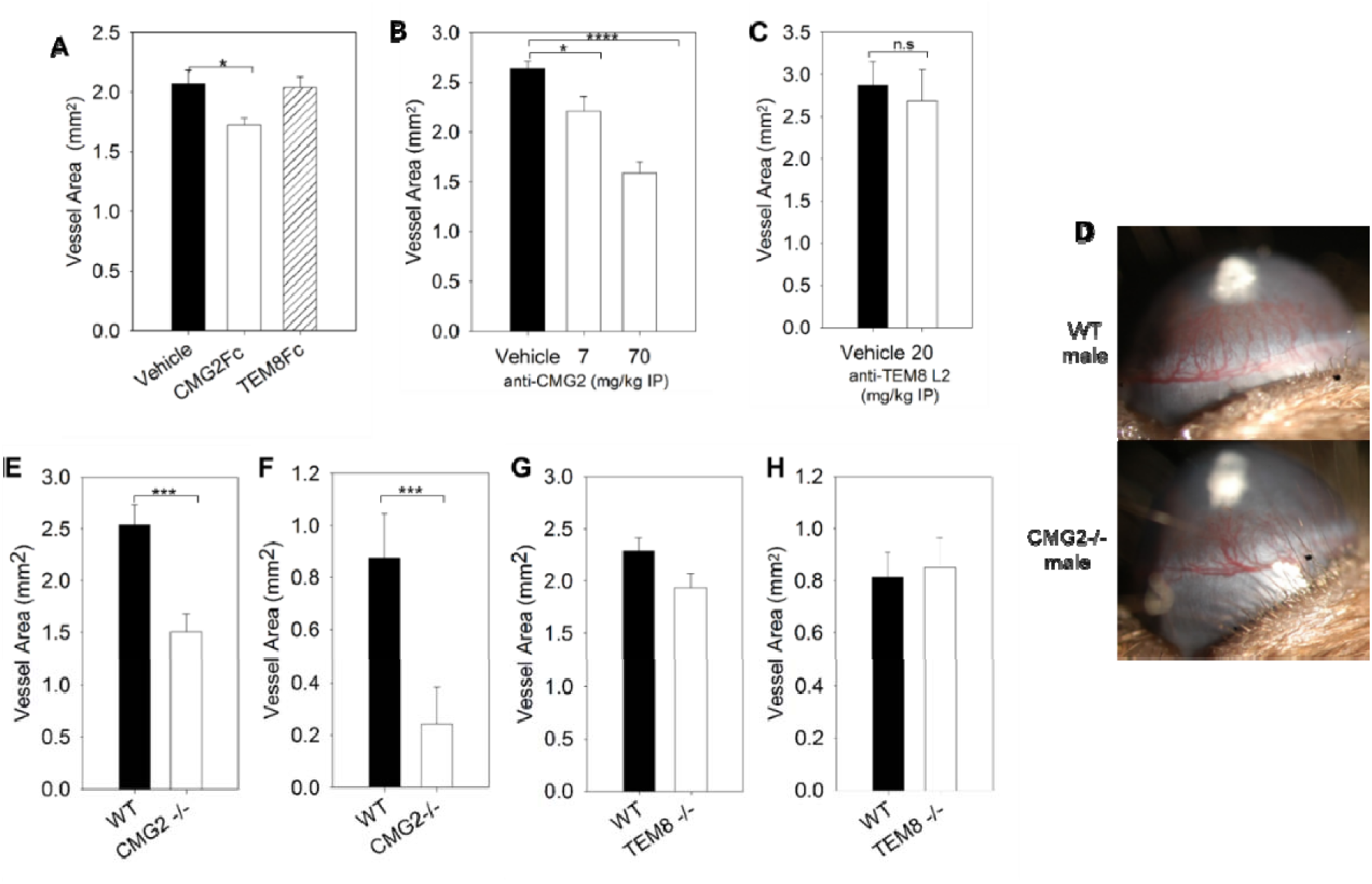
Blocking the interactions of CMG2 with its natural ligand inhibits both FGF and VEGF induced corneal neovascularization *in vivo.* (A) Corneal micropocket assay on mice treated with soluble ECD of either CMG2 or TEM8, via intraperitoneal injection. (B-C) Corneal micro-pocket assay on mice treated with either anti-CMG2 antibody (B) or anti-TEM8 L2 anti-body (C). Corneal neovascularization in these assays was induced by bFGF before treatment; vessel area on both the left and right corneas were measured from each mouse. (D) Representative image of corneal neovascularization in both wild type (WT) and CMG2 -/- male mice. (E-F) Comparison of corneal neovascularization between WT and CMG2-/- mice induced by either bFGF (E) or VEGF (F). CMG2-/- mice showed significant reductions in neovascularization for both bFGF-and VEGF-induced angiogenesis. (G-H) A similar experimen was performed with WT and TEM8-/- mice with either bFGF (G) or VEGF (H). Data presented are pooled from both genders. Error bars are standard error of mean, * p<0.05; ** p<0.01; *** p<0.001.

To confirm the importance of CMG2 rather than TEM8 as a mediator of angiogenesis in the cornea, we next sought to identify phenotypic changes that result from CMG2 or TEM8 knockout. We performed the corneal micropocket assay in both WT and CMG2 knockout (-/-), mice, using either bFGF or VEGF to induce vessel growth (Fig 1D-H). CMG2 knockout (CMG2-/-) mice exhibit a striking reduction in both bFGF-induced (Fig 1E) and VEGF-induced (Fig 1F) corneal vascularization compared to WT mice. This result clearly indicates a role for CMG2 in corneal angiogenesis. Interestingly, female mice were particularly susceptible to this effect (Fig S1A-B), and exhibited a dramatic (>85%) reduction in their response to VEGF (Fig S1B). In contrast, TEM8 knockout (TEM8-/-) mice exhibited no significant reduction when stimulated with VEGF (Fig 1H) and only modest reductions in bFGF-induced neovascularization (15%; p<0.05), (Fig 1G, S1C-D). These data confirm that CMG2 plays a quantitatively substantial role in corneal neovascularization, while TEM8 does not.

It remained possible that both anthrax toxin receptors might still be responsible for mediating the previously observed anti-angiogenic effects of PA^SSSR^. To assess this possibility, we evaluated the efficacy of PA^SSSR^ in reducing corneal vascularization in CMG2-/- or TEM8-/- mice. Importantly, CMG2-/- mice did not show reduction in bFGF-induced corneal angiogenesis (Fig2A). In contrast, PA^SSSR^ treatment significantly reduced corneal neovascularization in both wild-type control animals and TEM8-/- mice (Fig 2B). We conclude that CMG2 blockade is responsible for the anti-angiogenic effects of PA^SSSR^.

**Figure 2.**
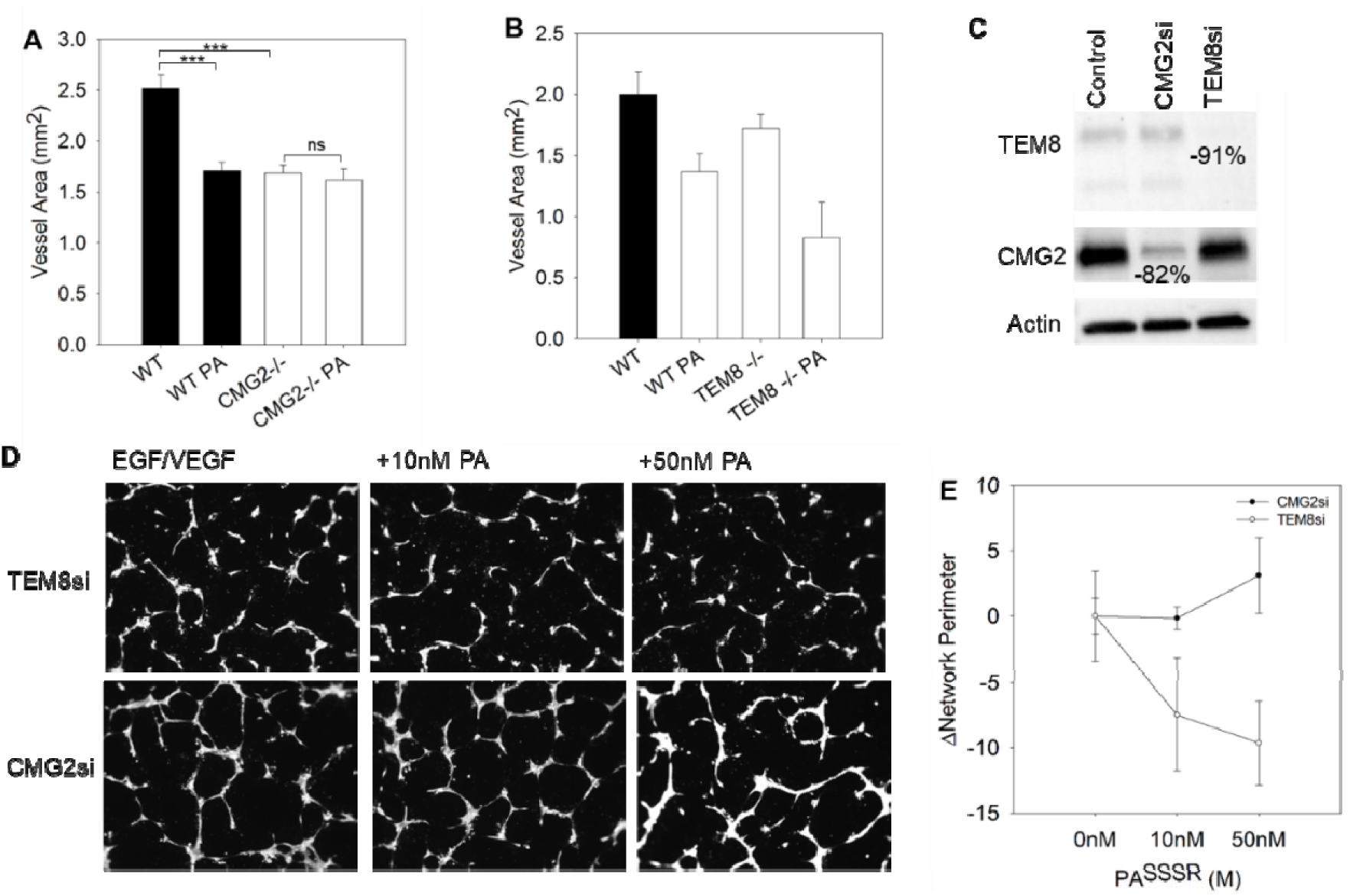
CMG2 is the main mediating receptor of PA^SSSR^-induced angiogenic inhibition. (A) Comparative levels of bFGF-induced corneal neovascularization between CMG2-/- and WT mice treated with or without PA^SSSR^, and (B) TEM8-/- and WT mice treated with or without PA^SSSR^. Results showed that PA^SSSR^ reduced vessel formation on TEM8-/- mice but not CMG2-/- mice. Error bars are standard error of mean. (C) Western blot of HMVEC lysates with either CMG2 knockdown (CMG2si) or TEM8 knockdown (TEM8si). Band intensity was normalized to control (untreated cells). (D) EGF- and VEGF-induced tubule formation assays with both CMG2si- and TEM8si-HMVEC, treated with or without different concentrations of PA^SSSR^. (E) Quantification of tube formation assays by counting the number of networks on each field. Error bars are standard deviation.

### Comparison of roles for CMG2 and TEM8 in endothelial tube formation, cell proliferation, and cell migration

To establish whether the effect of CMG2 knockout or blockade is intrinsic to endothelial cells, we performed tube formation assays in Matrigel with HMVEC cells, which express both CMG2 and TEM8. We note that due to culture limitations, both CMG2 and TEM8 ablation in HMVEC using CRISPR is not possible. However, knockdown using siRNA was successful. We compared the anti-angiogenic effects of PA^SSSR^ on cells selectively expressing only one of the two receptors (Fig 2C). In TEM8 knockdown (KD) cells (which primarily express CMG2), PA^SSSR^ administration resulted in a concentration-dependent reduction in the extent of tube network formation (Fig 2D-E). In contrast, tube formation in CMG2 KD cells was not altered by PA treatment (Fig 2D-E). Together with knockout data described above, these results demonstrate that PA^SSSR^ retains its anti-angiogenic effect in both TEM8-/- mice and TEM8 KD HMVECs but not in CMG2-/- mice or CMG2 KD cells. Hence, we conclude that PA^SSSR^ exerts its anti-angiogenic effects on endothelial cells via CMG2.

We next worked to determine the CMG2-mediated endothelial process that PA^SSSR^ disrupts to decrease angiogenesis. During angiogenesis, endothelial tip cells receive migratory signals and orient themselves to migrate up a chemotactic gradient, while stalk cells follow and then proliferate to form new vessel(s).^24^ Thus, CMG2 could regulate angiogenesis by impacting cell proliferation and/or cell migration. We have previously demonstrated that PA^SSSR^ affects cell migration, but not proliferation in HMVEC^1^, a primary endothelial cell type. Consistent with this result, CMG2 knockdown did not significantly reduce HMVEC proliferation (Fig S2A-D). Similar results are obtained for TEM8 knockdown. We also observe no reduction in cell proliferation in EA.hy926 endothelial cells (a fusion of human umbilical vein cells with lung carcinoma cells) treated with PA^SSSR^ under conditions that selectively target CMG2 (Fig S4A). Together, these data suggest that CMG2 inhibition must impact angiogenesis by modulating endothelial cell migration, rather than cell proliferation.

### Investigation of possible ECM ligands for CMG2

Our data support the idea that PA^SSSR^ exerts its anti-angiogenic effects by competing with interaction(s) between the CMG2 von Willebrand Factor A (vWA)^25, 26^ domain (including the metal ion-dependent adhesion site (MIDAS)^5, 27^ and endogenous ligand(s). However, the specific ligands that regulate angiogenesis by binding CMG2 remain unidentified, although different possibilities have been proposed^2, 8^. ECM proteins are likely candidates because of CMG2’s homology with integrins^26^, modest available data showing CMG2 interaction with ECM proteins^2^, and the observation that in individuals with hyaline fibromatosis syndrome (HFS), CMG2 mutations in the VWA domain result in widespread accumulation and of extracellular matrix (ECM) proteins^8^. To verify interaction of CMG2 with ECM proteins, we used ELISA to measure CMG2 K_d_ for a series of different ECM proteins (Table S1; Fig S3 A-D). Interestingly, each of the ECM proteins tested (Collagens I, VI, laminin and fibronectin) interacted with CMG2 with near-indistinguishable near-micromolar K_d_ values (Table S1). Similar *K*_d_ values for different ECM proteins in these assays did not reflect artefacts in the binding assay, since both positive and negative control assays (CMG2 + PA^SSSR^ and CMG2 + PA^SSSR^ in the presence of EDTA, respectively) mirrored previously published K values^18^. Previously published data on CMG2– matrix interactions used single-concentration assessment to conclude that CMG2 may bind preferentially to collagens IV^2^ or VI^8^, but did not measure relative CMG2 affinity for different ECM proteins. Differences in protein preparation, matrix adhesion to wells, number of binding sites per molecule, and incubation time are all factors which influence relative signal observed in blot- and ELISA-based assays, our binding curves are not necessarily at odds with previously reported results. Differences in protein preparation, matrix adhesion to wells, and number of binding sites per molecule are all factors which influence relative signal observed in blot- and ELISA-based assays, and our binding curves are not necessarily at odds with previously reported results^2, 8^. In addition, as a strategy to correct for possible differences in binding kinetics between ECM proteins, our ELISA data was quantified after fluorescence signal had largely stopped changing. Our initial time points showed much higher fluorescent signal for Col-VI than other ECM proteins, consistent with the single-concentration observations reported by the van der Goot lab^8^; however, the observed higher fluorescent signal was not indicative of higher affinity. Similar affinity of CMG2 for the matrix proteins tested, as shown in Table S1, indicates that CMG2 may bind to multiple ECM ligands and/or ECM peptide fragments. Such broad binding of ECM proteins also indicates that CMG2 could, like integrins, a significant role in cell adhesion and motility through direct interactions with the extracellular matrix.

### CMG2 and its role in cell adhesion and migration

To provide evidence for the assertion that CMG2 could play a role in cell adhesion, we performed *ex vivo* cell adhesion assays with EA.hy926 cells on plates coated with different ECMs (Collagens I, IV, and VI, Human Fibronectin, and Laminin-111). Treatment with PA^SSSR^ at concentrations identical to the CMG2 K_d_ value (200pM) significantly reduced EA.hy926 cell adhesion on plates coated with each ECM, but not on uncoated plates (NT) (Fig S4B). We note that at this concentration CMG2, but not TEM8, is blocked by PA^SSSR^. Together, these data suggest that CMG2 plays a role in mediating cell adhesion to ECM proteins^26^, and that PA^SSSR^ binding to CMG2 inhibits that interaction.

In angiogenesis a major function of cell adhesion is to enable cell migration. We hypothesize that CMG2 may play a role in enabling cell movement. In that case, PA^SSSR^ would inhibit cell migration as well as adhesion. Indeed, we find that PA^SSSR^ significantly reduces cell migration on multiple different ECM substrates (Fig S4C-D). Specifically, we used fetal bovine serum as chemoattractant in a microfluidic perfusion device which contains a perfusion chamber that allows formation of a small molecule- and macromolecule-concentration gradient. In this setup, we can observe migration of individual cells in real time. As expected, PA^SSSR^ treatment resulted in decreased migration over nearly all ECM surfaces (Fig S4C-D), even at the 200pM concentration expected to result in only 50% bound CMG2. Combining these adhesion and migration data, we conclude that ECM proteins mediate endothelial cell adhesion and migration through interaction with CMG2, and that PA^SSSR^ can disrupt this interaction (Fig S4B-D).

Notably, the nature of this migration assay allows us to compare two different aspects of migration: initiation of cell movement (chemokinesis) and directional motion towards growth factor-containing serum (chemotaxis). Surprisingly, targeting CMG2 in EA.hy926 cells with PA^SSSR^ at the CMG2 Kd (200pM) showed a 70% decrease in observed chemotaxis on non-coated surface (70%, P<0.01; Fig S4C. Given that we expect only 50% CMG2 occupancy at these PA^SSSR^ concentrations, this chemotactic inhibition is dramatic and surprising. In contrast, the total distance traveled by PA^SSSR^ treated cells was only modestly (∼20%) reduced, compared to untreated cells (Fig S4D). Thus, CMG2 targeting has a dramatic effect on chemotaxis but little effect on chemokinesis.

Futher support for involvement of CMG2 in chemotaxis rather than chemokinesis is provided by transwell migration experiments performed on CMG2 knockdown HMVEC cells. Like many other traditional migration assays, the transwell assay does not provide a stable chemoattractant gradient to allow chemotaxis measurement^28^, and thus only measures chemokinesis. Evaluation of these CMG2 knockdown cells via transwell migration assays showed no significant difference between wild-type and CMG2 or TEM8 knockdown HMVEC (Fig S4E), consistent with our previous observation that CMG2 inhibition impacts chemotaxis but not chemokinesis.

We next examined the effect of CMG2 depletion on chemotactic migration in EA.hy926 cells, using CRISPR to target Exon1. Loss of CMG2 function in knockout cells was validated with a flow cytometry assay in which uptake of a fluorescent PA-AF568 conjugate was evaluated at CMG2 K_d_ 200pM (Fig 3A). CMG2 knockout cells showed substantial inhibition of intracellular fluorescence relative to WT cells, and knockout cell fluorescence was indistinguishable from that of negative control WT cells (Fig 3A). The ability of transfected CMG2-mClover to rescue the PA uptake defect was confirmed, demonstrating that off-target effects cannot explain loss of PA uptake in these cells (Fig 3A). Importantly, evaluation of the migration of these EA.hy926 knockout cells in our microfluidic device demonstrated that while WT cells migrated toward serum (Fig 3B), CMG2-/- cells migrated only in a random manner (Fig 3C). Significantly, knockout cells showed complete abolition of chemotactic migration (Fig 3E) but only slight alteration of chemokinetic migration (P<0.001) (Fig 3F).

**Figure 3.**
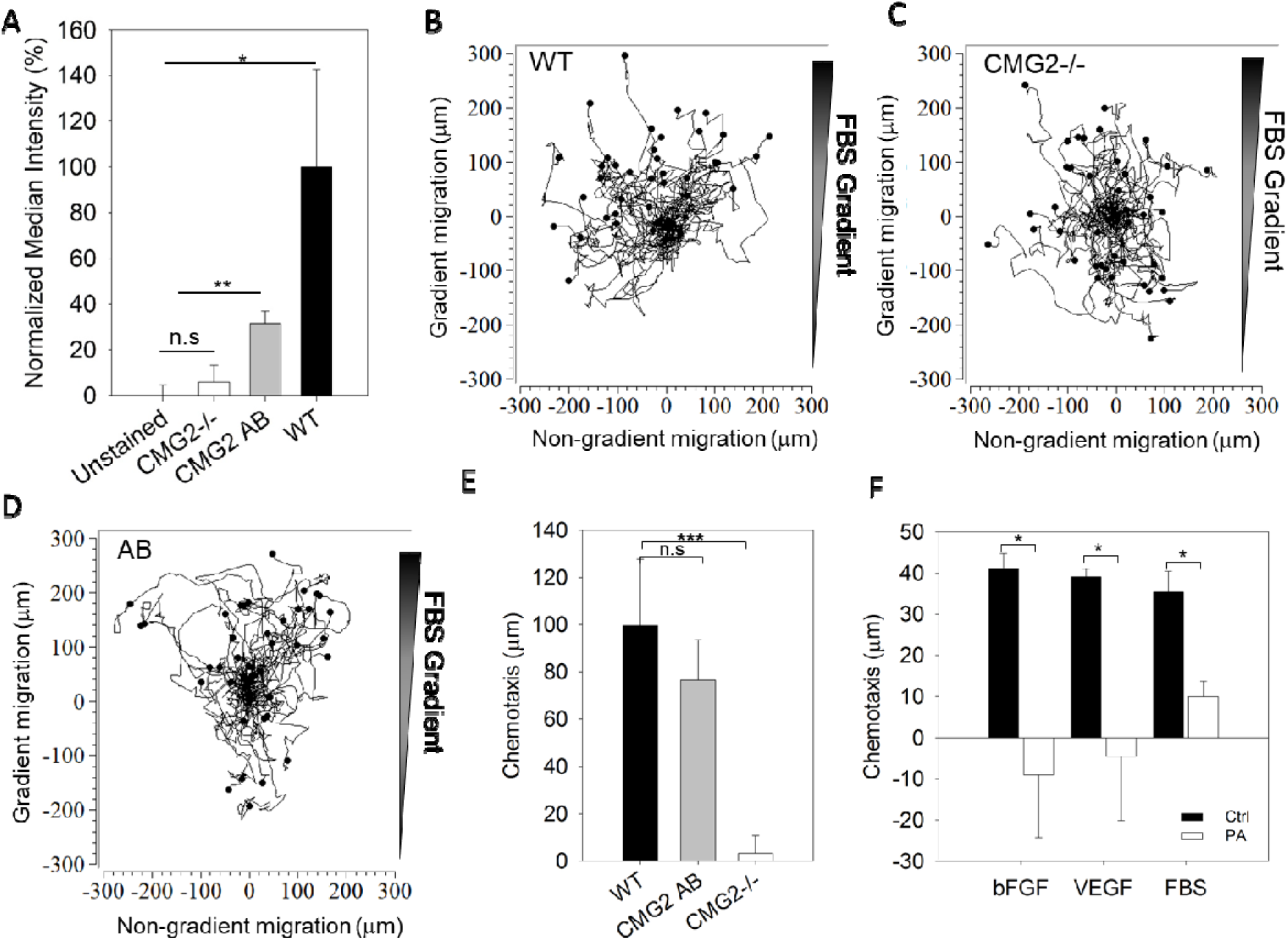
CMG2 KO EA.hy926 endothelial cells lose the ability to migrate toward chemoattractants. (A) Differential PA-AF586 conjugate uptake by wild type (WT), CMG2 add back (AB) CMG2 knockout (CMG2 -/-) EA.hy926 cells via flow cytometry (10,000 cells per condition). Median fluorescence intensiy of each condition was normalized against WT signal after subtracting from the unstain control. Error bars are normalized standard deviation from three individual replicates. (B) Aggregated track plots of individual EA.hy926 cell migration in the CellASIC migration chamber. Both WT cells (B), CMG2-/- cells (C), and CMG2 AB cells were subjected to an FBS gradient. (E) Quantification of EA.hy926 chemotaxis (displacement toward gradient, E) in the CellASIC migration assay. (F) Quantification of EA.hy926 WT cells chemotaxis in bFGF, VEGF and FBS with or without the presence of 200pM PA^SSSR^. * p<0.05; ** p<0.01; *** p<0.001; n.s not significant

To confirm that the loss of chemotaxis in CMG2 KO cells was not a consequence of off target CRISPR effects, we complemented the CMG2 mutation with a CMG2-clover expression vector and found that CMG2 add-back restored EA.hy926 cell chemotaxis (Fig 3D-E). Hence, CMG2 is a critical component of chemotactic sensing or response in these cells, and may play a role in regulating endothelial cell chemotaxis *in vivo*. Of interest, CMG2 is required for reorientation of the mitotic spindle in zebrafish embryos^29^ suggesting that it might also play a role in orienting the microtubule cytoskeleton in a chemotactic context.

### CMG2 impacts developmental angiogenesis

We next investigated the effect of CMG2 deletion on developmental angiogenesis. To do so, we compared retinal vessel development between WT and CMG2-/- mouse. We found that the total vascular area was similar between the two genotypes (Fig. 4). However, the vascular growth pattern observed was different between knockout and WT mice, with knockout mice having a more bush-like pattern versus the tree-like pattern observed in WT animals (Fig. 4A). Careful quantitation showed that the total number of veins in the retina of WT and CMG2-/- mice is similar (Fig 4A-B), but CMG2-/- mice have slightly fewer primary arteries (Fig 4A, C). Importantly, CMG2-/- mice showed 2.5-fold more primary branches per artery than the WT cells (Fig 4A, D). This difference in vessel patterning could be consistent with the loss of chemotactic migration observed in knockout cells. We speculate that in the absence of CMG2, endothelial cells effectively lose chemotactic signals (though haptotactic and other signals may be retained), which reduces effective directional movement. This (hypothesized) weaker response in CMG2-/- mice to a developmental angiogenic gradient would leave some areas of the retina with poor perfusion. Angiogenic signals from these areas could then lead to additional branching to alleviate hypoxia, resulting in a vasculature that remains functional, but is patterned differently.

**Figure 4.**
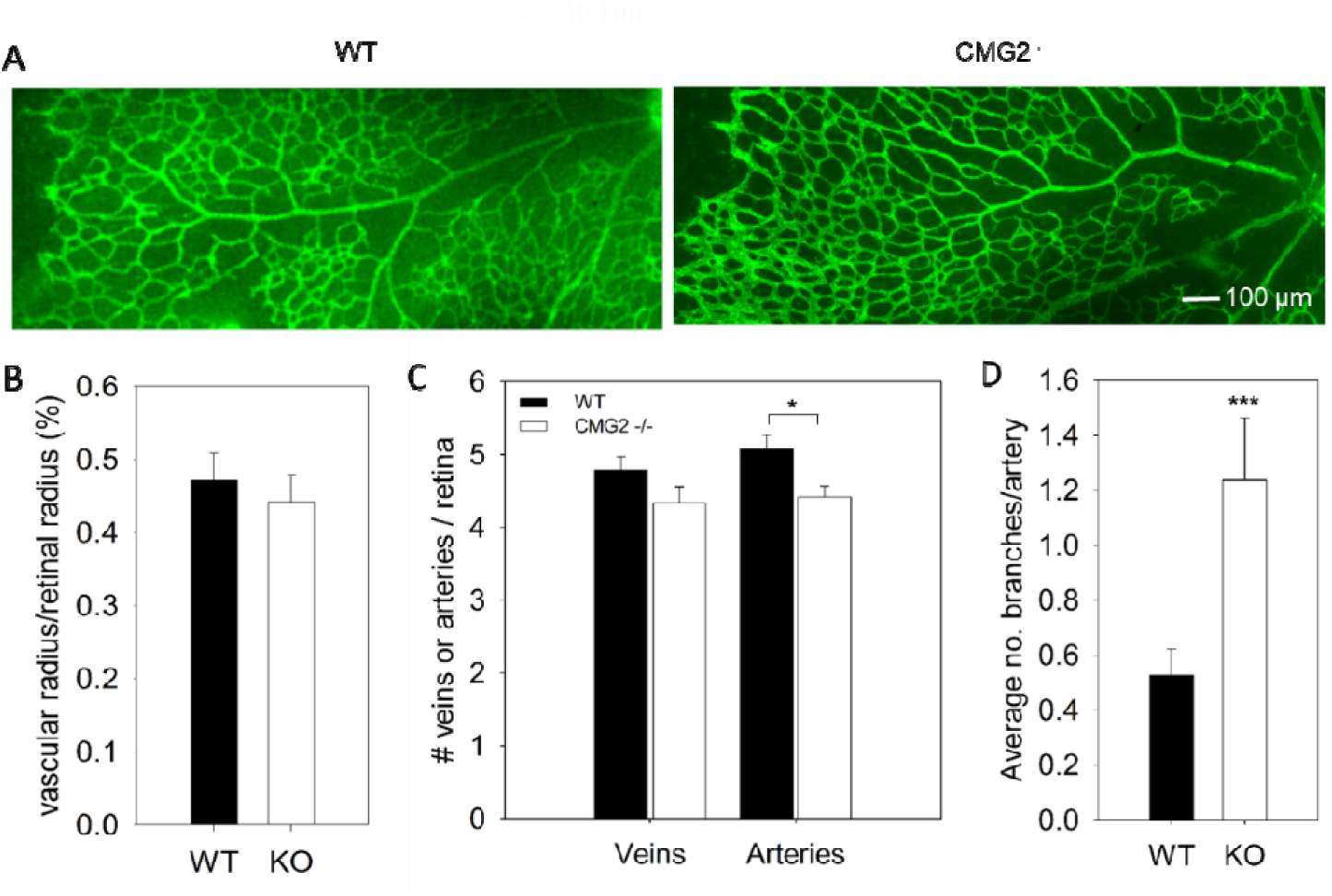
CMG2 KO increases vessel branching in the mouse retina. (A) Representative mages of vessel formation in the retina of both WT (left) and CMG2-/- (right) mice. (B-D) Quantified vessel formation from retinal assays. (B) Comparison of vascular radius, normalized to the retinal radius, between WT and CMG2-/- mice. (C) Quantification of arterial branching in both WT and CMG2-/- mouse retinas. (D) Artery and vein counts per retina as in WT and CMG2-/- mice. CMG2-/- mouse retinas exhibit fewer veins and arteries than WT, but only artery count is significantly lower than WT mice.

### CMG2-mediated chemotaxis is sensitive to multiple growth factors

We have demonstrated here that CMG2-deficient mice showed potent reduction in corneal neovascularization induced by both bFGF and VEGF. To more fully elucidate CMG2’s angioenic response to bFGF and VEGF, we carried out experiments that measured endothelial cell chemotactic migration in response to these two growth factor gradients respectively. EA.hy926 cells display chemotactic migration towards FBS, bFGF and VEGF (Fig S5), and this migration is significantly inhibited with 200pM of PA^SSSR^ treatment (Fig 3F, S5). Cells treated with PA^SSSR^ showed significantly lower chemotaxis in FBS, bFGF, or VEGF gradient (Fig 3F, S5), further suggest that CMG2 plays a critical role in mediating chemotaxis towards multiple growth factors that are in the serum. Together, these data suggest that CMG2 may act as a central regulator for chemotaxis that is initiated by bFGF and VEGF. The mechanism of CMG2’s impact on both bFGF and VEGF induced chemotaxis will be important for developing new efficacious therapies to treat pathological angiogenesis. It remains to be seen whether interaction of CMG2 with growth factors receptors at the cell surface can be detected.

## Methods

### Protein preparation

#### Cell culture

EA.hy926 (CRL-2922) cells are the result of a fusion of human umbilical vein cells with lung carcinoma cells. Cells were cultured in 10% FBS + DMEM and incubated at 37°C in a humidified environment with 5% CO_2_ until ready for passaging.

#### EA.hy cell proliferation assay

15,000 EA.hy926 cells were seeded into each well in a 96 well plate and incubated for 1 h to attach. After cell attachment, media with treatments were added. Ethanol fixed cells were used as negative control. After a 24 h incubation, 20 µL CellTiter-Blue Reagent (Promega) was added to each well for 4 h. Fluorescence signal (Ex: 560nm / Em: 590nm) was measured using a BioTek Synergy H2 plate reader. All readings were normalized to the non-treated control.

#### Cell ASIC migration assay

The assay protocol followed the CellASIC ONIX M04G-02 Microfluidic Gradient Plate User Guide. All media put into the plate (excepting the cell suspension) was filtered through a 0.2 µm syringe filter. EA.hy cells (3 x 10^6^ cells/mL) were loaded in and incubated overnight with DMEM + 10% FBS under flow at 37°C. Assays were then performed with a stable gradient of DMEM + 0 – 10% FBS with or without peptide treatment. Brightfield images at 10x magnification were taken every 10 min over 12 h on an Olympus IX73 microscope and the ORCA-Flash4.0 camera (Hamamatsu). Individual cells were tracked with Image J manual tracking plugin. Data was transferred into the ibidi Chemotaxis and Migration Tools 2.0. to export endpoint y displacement and accumulated displacement. P-values were calculated using Student’s *t*-test and error bars are standard error of the mean (n=50).

#### CMG2 ECM ELISA

All steps were performed at room temperature unless otherwise indicated. For binding assays, matrix protein (Rockland Immunochemicals, Corning, EMD Millipore) was adsorbed onto 96-well polyethylene plates (Greiner) by incubating 2 ug/mL matrix protein in HBS with 2 mM Mg^2+^ and Ca^2+^ (Buffer A) in individual wells at 4° C overnight. Bacillus anthracis protective antigen (PA) was treated similarly but was incubated at 1 μM in the same buffer. After incubation, wells were washed 3x with Buffer A and blocked with 5% BSA (GoldBio) in HBS with 2 mM Mg^2+^, 2 mM Ca^2+^, and 0.1% Tween-20 (Buffer B) for 1 hour. Blocking was followed by 3 washes with Buffer B, after which varying concentrations of a biotinylated CMG2-AviTag construct (for matrix, 2138 to 10 μM; for PA, 4 μM to 1 pM) were dissolved in Buffer B and incubated in wells for 4 hours. After incubation with CMG2-AviTag, wells were again washed 3 times, after which streptavidin-HRP (Thermo Scientific) diluted in Buffer B with 5% BSA was incubated in wells for 1 hour. Wells were then washed 6 times with Buffer B, after which 1x TMB solution (Thermo Scientific) was added to wells. Once color was visible in wells, reaction was quenched with 0.2 M H_2_SO_4_. Wells were read out in a BioTek H4 Hybrid plate reader (BioTek) by quantifying well absorbance at 450 nm. Data were analyzed in Microsoft Excel, and Kd values were calculated in MATLAB using the sigm_fit function.

#### Adhesion assay

EA.hy 926 cells were grown in DMEM supplemented with 10% FBS. 20μg/mL collagen I, IV, VI, laminin, fibronectin, PA^SSSR^, and BSA were preared in PBS and coated in 96well plate (100Lg/well) overnight at 4°C. After coating, collagen I, IV, VI, laminin, fibronectin, PA^SSSR^, and BSA were removed and 5% BSA were coated for 1 hour. Cells were deprived with serum by scraped and cells were at 100g for 10 minutes and the supernatant was discarded. Cells then resuspended at 2-3×10^5^ cells/mL in DMEM. 100μL of cells were added to each of the coated wells. The plate was incubated at 37°C for 1 hour to allow the cells to adhere on the surface. Each well was washed gently for two times using warm serum-free DMEM by multi-channel pipettor. 100μL of serum free DMEM and 20μL of cell-titler blue (Promega Cat#G8080) were added into each well, and incubated for 4 hours at 37°C. After 4 hours incubation, fluorescent signal of 96 well plate was measured at 560_Ex_/590_Em_ by a plate reader.

#### EAhy 926 CMG2 KO development

HEK 293T cells (3.8 x 10^6^ cells) were seeded in 10cm tissue culture dish for 18 hours. 12μg of pCMV, 5μg of pVSVG, 12.5μg of Lenti-CRISPR (targeting CMG2 exon 1, sgRNA sequence: GCACCAACAGCCACAGCCCG), 90μL of 1mg/mL PEI and 600μL of serum-free DMEM were mixed into a tube and incubated for 15 minutes before adding to the 10cm culture dish. The dish was replaced with 10mL of 10% FBS DMEM in 4 hours and followed by 48 hours incubation. Media was removed from the transfected cells into a conical tube, and was spin at 2500g for 3 minutes to remove debris. The supernatant (lentivirus) and 10μg/mL polybrene were added to 40% confluent EA.hy926 cells and incubated for 24 hours. The next day, media was removed, fresh 10% FBS DMEM and 1μg/mL puromycin were added to the plate for 3-5 days. After 3-5 days, the cells that survived in the selection were diluted into single colony in 96 well plate.

#### PA uptake flow cytometry

CMG2 KO EA.hy 926 cells and WT EA.hy926 cells were split in 12 well plate at 50% confluent. 200pM PA-Alexa Fluor 546 was added into each well and incubated overnight. Then cells were trypsinized and resuspended into a microcentrifuge tube. Cells were washed once with complete media. Then 50,000 cells of each condition were sent to Cytoflex (RIC facility at BYU). Data was collected and analyzed in our lab by using FlowJo software.

#### Corneal micropocket assay

The corneal micropocket assay was performed as described^16^ using pellets containing 80 ng of basic fibroblast growth factor (bFGF) or 180 ng of human carrier-free recombinant vascular endothelial growth factor (VEGF; R&D Systems) in C57BL/6J mice. The treated groups received daily or twice daily i.p injections for 5 (bFGF) or 6 (VEGF) consecutive days of protein in PBS. Treatment was started on the day of pellet implantation; control mice received only vehicle i.p. The area of vascular response was assessed on the 5th (bFGF) or 6th (VEGF) postoperative day using a slit lamp. Typically, at least 10 eyes per group were measured.

#### HMVEC proliferation assay

Human microvascular endothelial cells (Cambrex) were maintained in EGM-2 (Cambrex) according to the vendor’s instructions and used before passage 7. On day 0, proliferating cultures of BCE or HMVEC-d cells were seeded at ∼ 10% confluence into 96-well plates. After attachment, medium was exchanged for medium containing 1 pmol/L to 1 μmol/L of the indicated protein. Cells were allowed to grow for 7 days and then quantified using CyQUANT (Invitrogen) according to manufacturer’s directions. The degree of proliferation in culture was measured by comparing wells in each plate fixed in absolute ethanol on day 0 with experimental wells, with fold proliferation calculated by dividing CyQUANT fluorescence in experimental wells by that in day 0 wells. Groups were compared using Student’s *t* test, with Bonferroni correction where appropriate.

#### HMVEC Trans-well migration assay

Human microvascular endothelial cells were maintained as above. Polycarbonate Transwell inserts, 6.5 mm diameter with 8.0 μm pores, were coated with fibronectin (BD Biosciences). Cells were harvested and resuspended in EBM (Cambrex) containing 0.1% bovine serum albumin (Fisher Chemical). Cells (10,000 per well) were plated onto wells containing medium alone or medium containing the molecule to be tested. These wells were suspended above wells containing 5 to 10 ng/mL recombinant human VEGF (R&D Systems) or full serum medium. Cells were allowed to migrate for 4 h. Membranes were rinsed once in PBS and then fixed and processed using Diff-Quick (Dade Diagnostics). Cells on the top of the membrane were removed using cotton-tipped applicators, and the membrane was removed from the insert using a scalpel. Membranes were then mounted on slides, and the number of cells in a microscopic field was counted manually.

#### HMVEC CMG2 / TEM8 KD

CMG2 and/or TEM8 knockdown in HMVEC was achieved using Dharmacon smartpools (Dharmacon, Lafayette, CO) and RNAimax (ThermoFisher, Waltham, MA) in OPTI-MEM with Glutamax (ThermoFisher, Waltham, MA). Approximately 6 hours after addition of the transfection complex, an equal volume of EBM-2 was added to the cells, and media was changed to EBM-2 ∼18 hours later.

#### HMVEC tube formation assay

Human microvascular endothelial cells were maintained as above. Before the assay, a 1- to 2-mm layer of Matrigel was plated into the wells of a 12-well cluster. Approximately 10^5^ cells were plated on this layer in EGM-2. Plates were examined at 12, 14, 16, 18, and 24 h for differences in network formation. In each experiment, good network formation was observed in untreated control wells.

#### CMG2 and TEM8 KO mice

CMG2 and TEM8 knockout mice were a kind gift from Stephen Leppla. They were housed in the ARCH facility at Boston Children’s Hospital on standard food and bedding (CMG2) or on standard food and bedding, with supplemental soft food available (Nutra-gel mouse diet, Bio-Serv, Flemington, NJ). Genotyping was performed by Transnetyx (Cordova, TN).

#### Western Blotting

Near-confluent HMVEC WT and CMG2 / TEM8 KD cells were washed three times in cold PBS, and lysed for 30min at 4°C in PBS supplemented with 1% Triton X-100 and a protease inhibitor cocktail (cOmplete mini protease inhibitor cocktail, Roche). Lysates were cleared by centrifugation at 16,000 xg and the supernatant protein concentration was measured by BCA assay. Protein (40μg) from each sample was used for SDS-PAGE and blotted onto PVDF. Following blocking, the membranes were probed with antibodies against CMG2 (1:1000 16723-1-AP, Proteintech, Rosemont, IL) or TEM8 (1:1000 ab21269, Abcam, Cambridge, UK), then stripped and reprobed for beta-actin (1:10,000 A5441, Sigma) as loading control. Blots were imaged using a ChemiDoc imager (Bio-Rad, Hercules, CA).

---

**Supplementary Figure S1.**
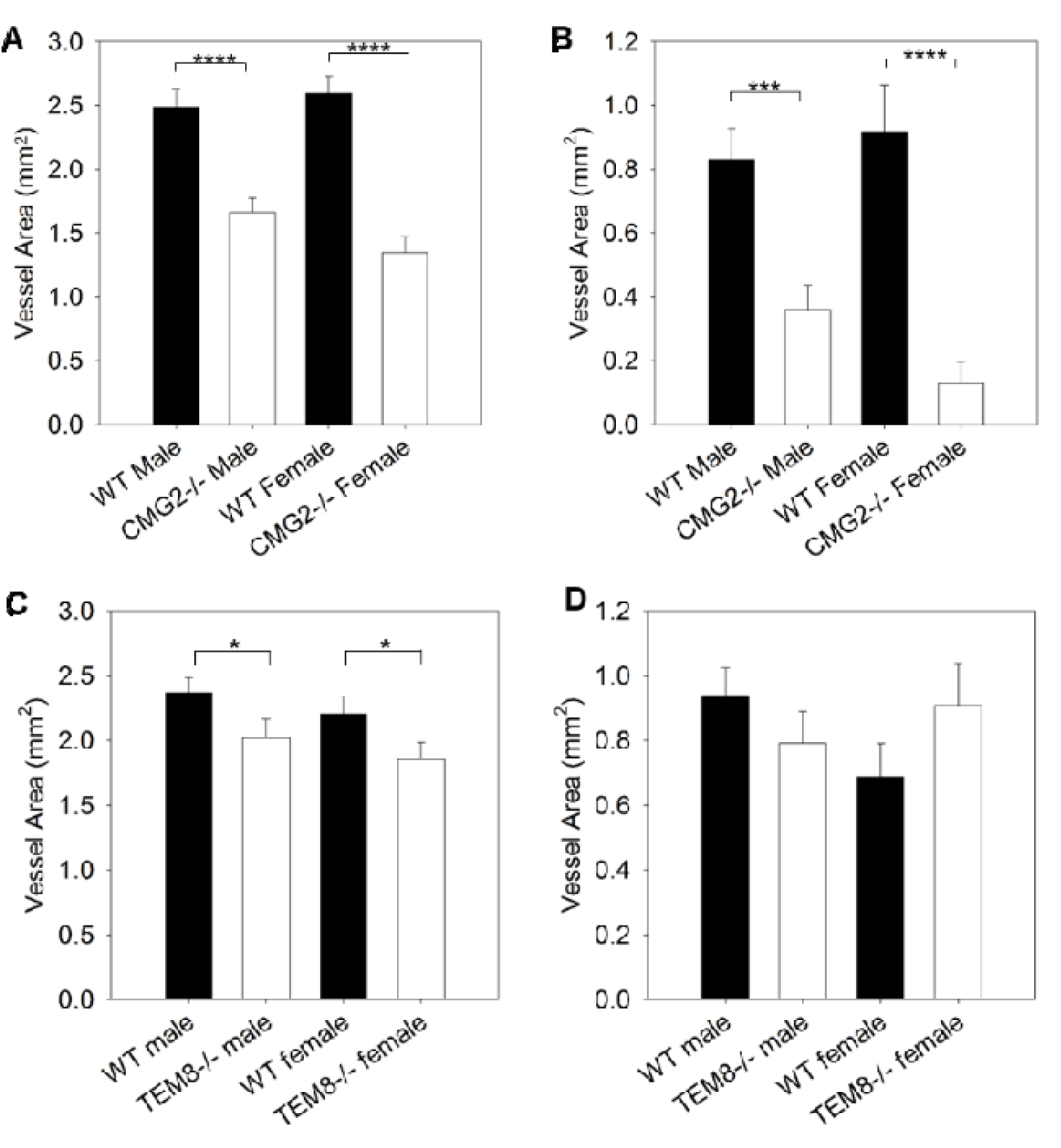
Gender-specific quantification of bFGF- and VEGF-induced corneal vascularization in WT, CMG2 KO, and TEM8 KO mice. (A-B) Quantification of (A) bFGF-induced and (B) VEGF-induced neovascularization in male and female CMG2-/- mice as compared to WT. (C-D) Quantification of (C) bFGF-induced and (D) VEGF-induced neovascularization in male and female TEM8-/- mice as compared to WT. For both genders, greater decreases in corneal vascularization were observed in CMG2 KO than for TEM8 KO.

**Supplementary Figure S2.**
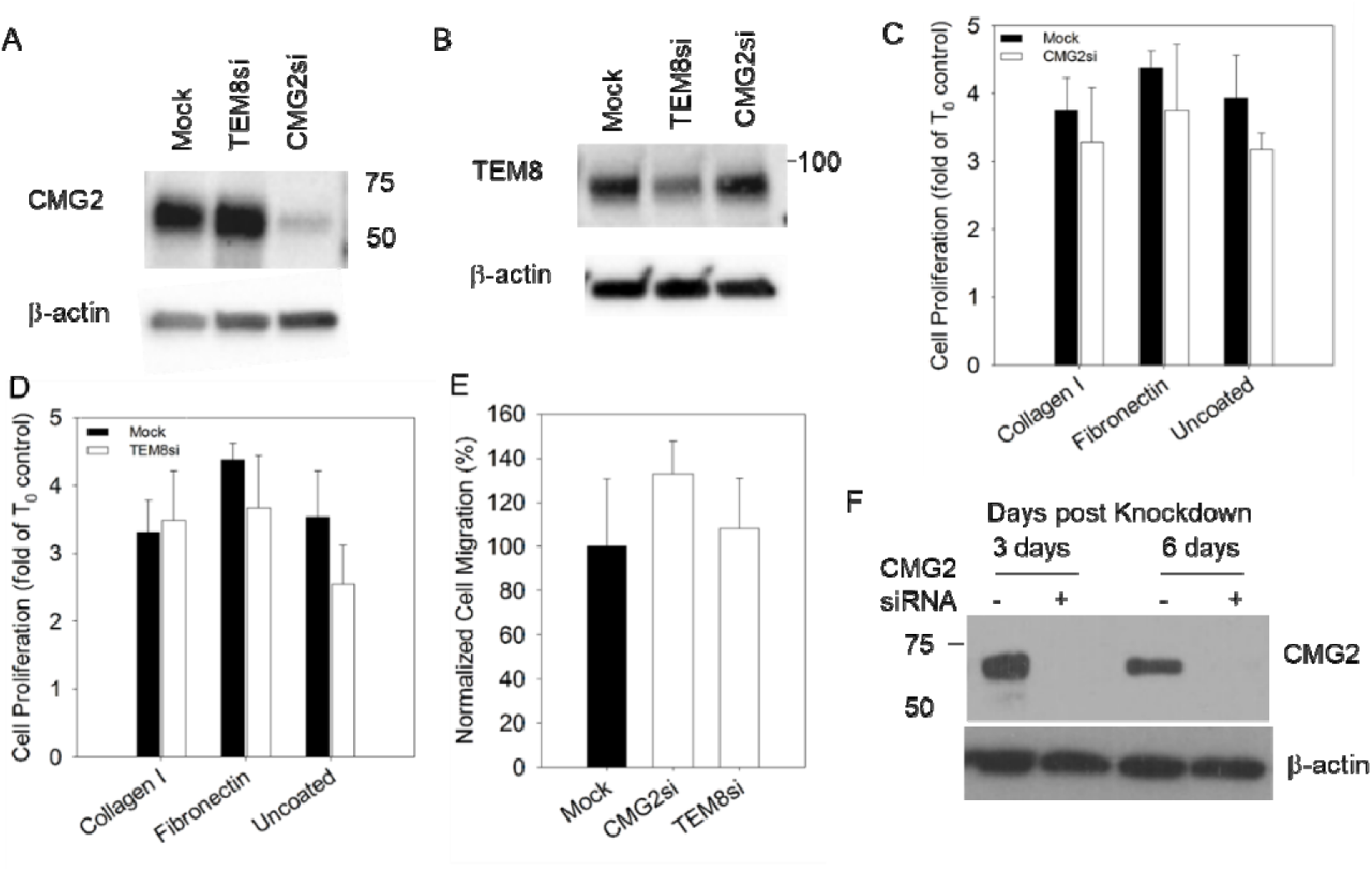
Effects of CMG2 and TEM8 siRNA-mediated knockdown on HMVECs cell proliferation and migration. (A-B) Western blot analysis of HMVECs lysates for (A) CMG2 and (B) TEM8 before and after introduction of siRNA. (C-D) Proliferation of both WT and siRNA-treated Ea.hy926 cells with both (C) CMG2-specific and (D) TEM8-specific siRNA. Proliferation was not significantly altered from WT in either CMG2-targeted or TEM8-targeted cells. (E) Migration of WT, CMG2 siRNA and TEM8 siRNA-treated HMVECs cells. No significant difference in migration was observed from WT. (F) Western blot analysis of HMVECs lysates for CMG2 expression 3 days and 6 days after addition of siRNA.

**Supplementary Table S1.** CMG2 binds several broadly expressed matrix proteins with high affinity. Results of an ELISA-based quantification of CMG2 binding to several broadly-expressed matrix proteins, including collagens, laminin-111, and fibronectin. For positive control, PA was assayed for binding to CMG2. Such similar affinities between matrix proteins indicate that CMG2 shows no preference for binding to any one of the matrix proteins assayed.

| Protein | kD (nM) | Std. Dev | Repeats |
| --- | --- | --- | --- |
| Collagen I | 500 | 141 | 3 |
| Collagen VI | 800 | 411 | 3 |
| Laminin-111 | 500 | 158 | 2 |
| Fibronectin | 500 | 141 | 3 |
| PA (positive control) | 0.46 | 0.047 | 3 |
| PA w/EDTA (neg. control) | None | N/A | 2 |

**Supplementary Figure S3.**
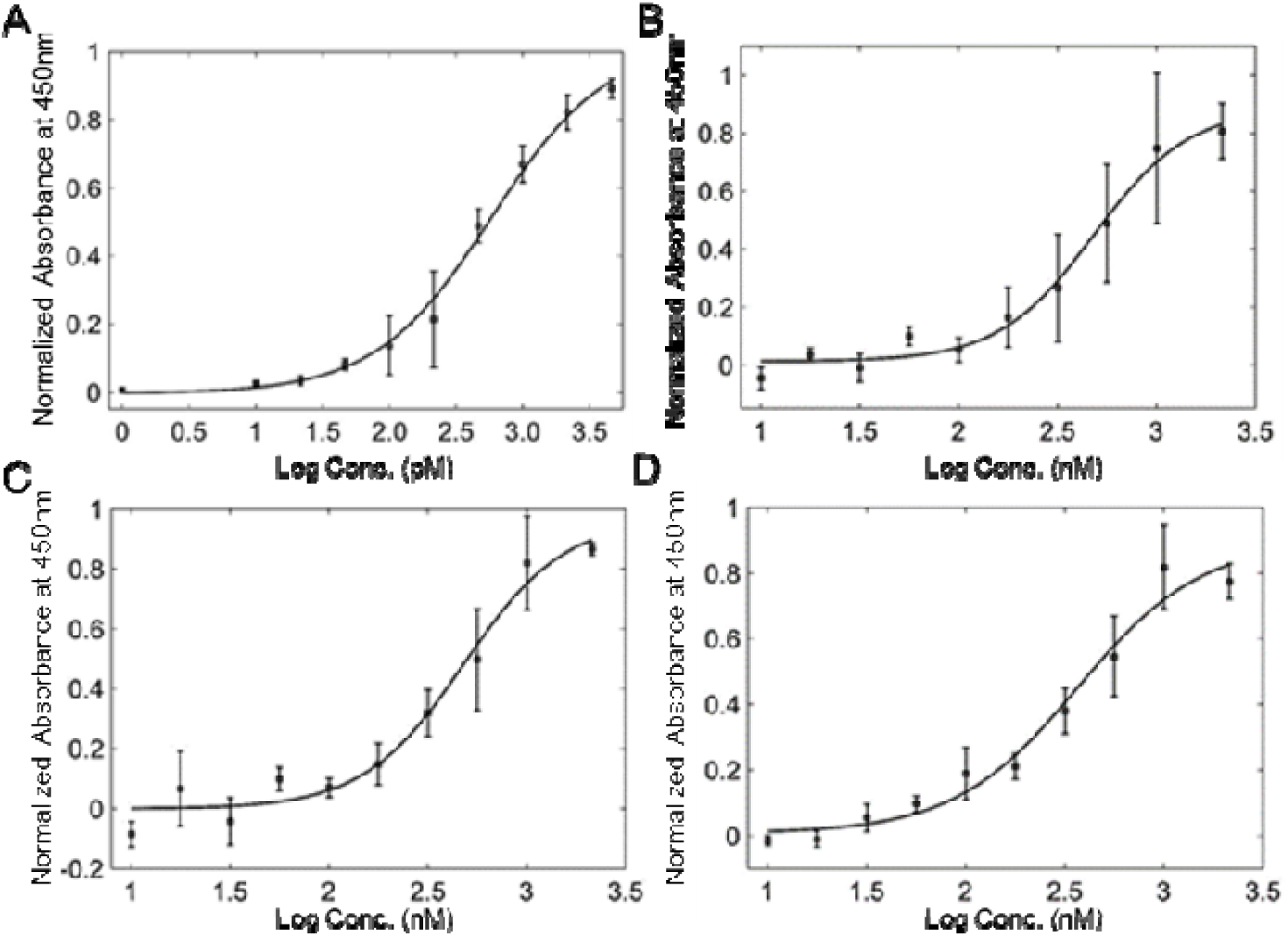
Binding of CMG2 to extracellular matrix proteins, as determined by ELISA. Proteins were coated on wells, after which CMG2-GST-biotin was added and signal read out with streptavidin-HRP/TMB. KD values were calculated by fitting each dataset to a 4-parameter logistic curve. A, positive control (PA); B, fibronectin; C, collagen I; D, collagen VI. PA x axis scale is logarithmic with pM concentration units; all others are logarithmic with nM concentration values. Y axis is normalized absorbance at 450 nm. Collagen IV and laminin-111 were also assayed for binding, but curves are not displayed due to poor fit. In each case, similar binding affinites (500 nM-1 uM) were observed (see Supplementary Table S1).

**Supplementary Figure S4.**
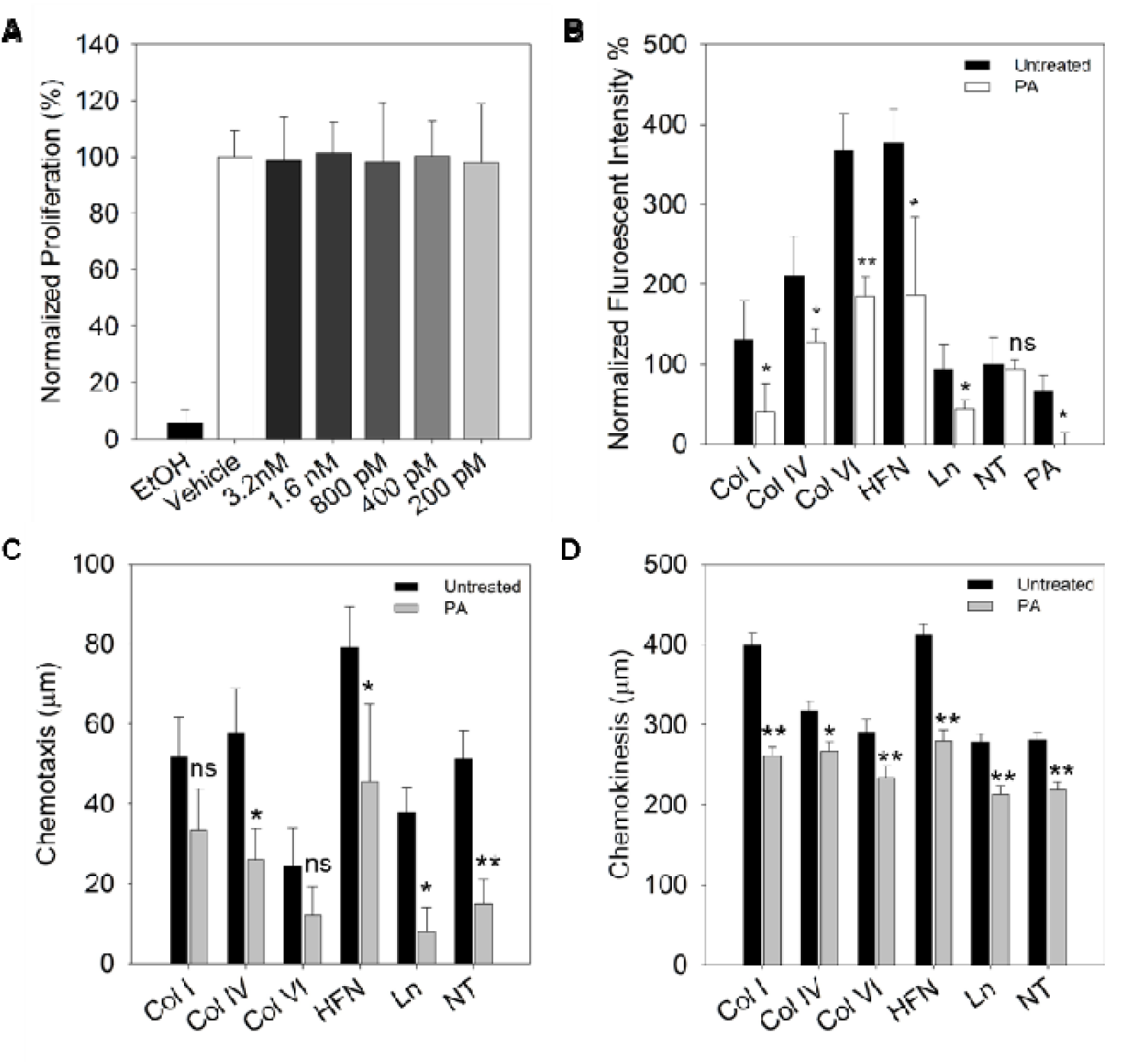
Matrix binding to CMG2 is an important component of endothelial cell adhesion and migration. (A) EA.hy926 endothelial cell proliferation in different concentrations of PA^SSSR^. Ethanol (EtOH) treated cells was used as the negative control. (B) EA.hy926 endothelial cell adhesion to plates coated with various matrix proteins, or no ECM proteins (“NT”), both with and without 200 pM PA. Addition of PA significantly inhibits adhesion to matrix-coated plates. Adhesion to uncoated plates was not affected by PA^SSSR^ treatment. (C-D) EA.hy926 migration on different ECM coated surface and compare to the PA^SSSR^ treatment. Chemotaxis are the vertical displacement measured towards a serum gradient form across the migration chamber (C). While chemokinesis are measured by total distance cell migrated in random direction (D). statistic significance between untreated and PA treated cell was calculated by student T.Test. Error bars are standard deviation of mean, * p<0.05; ** p<0.01; n.s not significant

**Supplementary Figure S5.**
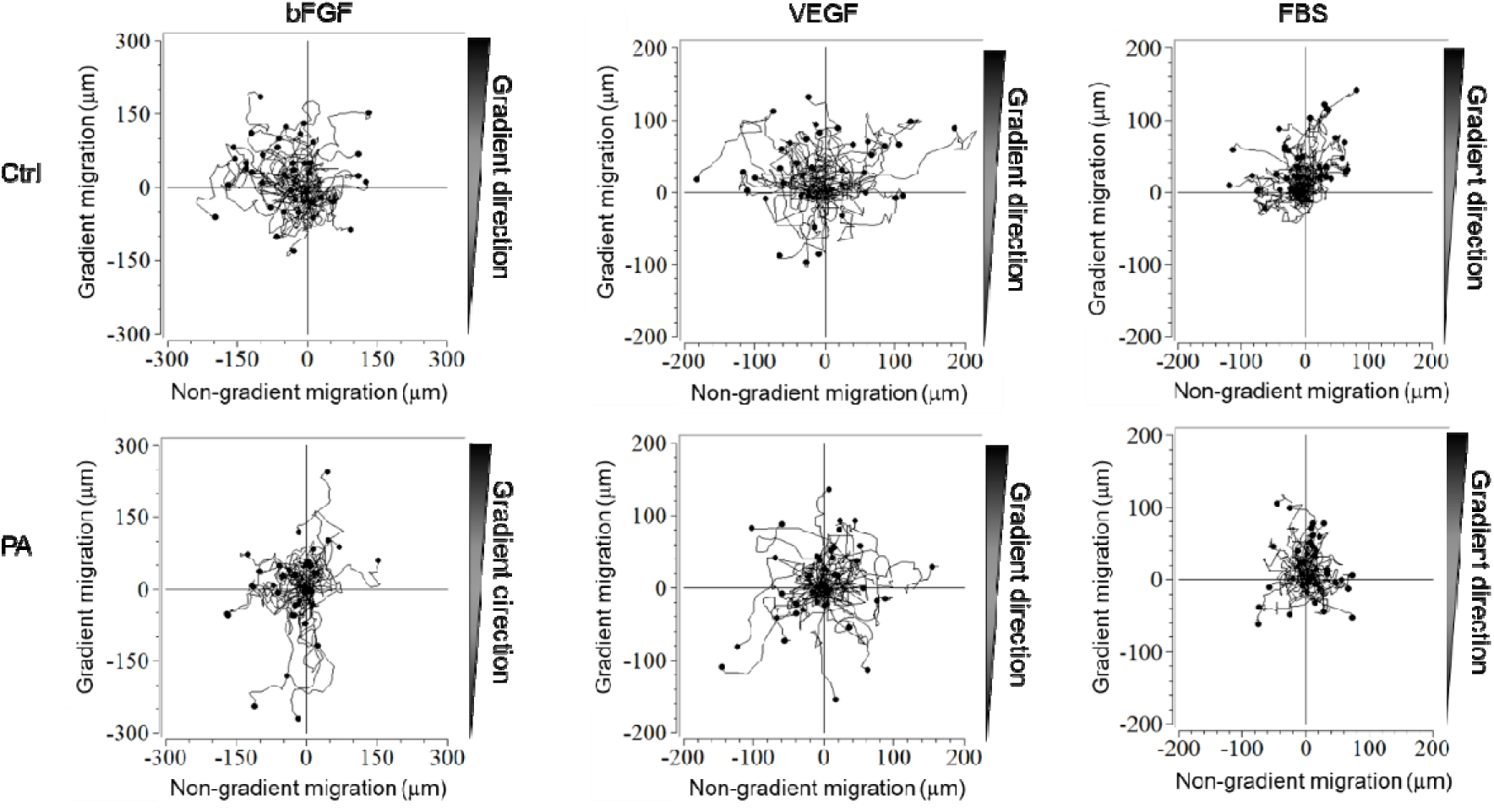
Migration track plots of CMG2 directs cell chemotaxis towards bFGF, VEGF, and serum. EA.hy926 WT cells were placed in a gradient of bFGF (left), VEGF (middle) and FBS (right), with or without the presence of 200pM PA^SSSR^ (top and bottom respectively). Each migration condition were performed 37°C for 8 hours. Cell tracking was done in ImageJ with forty cells were tracked in each condition. Raw data were then imported into chemotaxis and migration tool from ibidi to make track plots.

## Supplementary note

1. bFGF was more used in corneal assays because we saw a better angiogenic effects
2. 200pM PA was targeting CMG2

